# Proteomic profiling of xenobiotic and nutrient transporters in human placenta of different gestational ages

**DOI:** 10.64898/2026.06.25.730994

**Authors:** Eric M. Weaver, Ariel Topletz-Erickson, Nina Isoherranen, Jashvant D. Unadkat, Samuel L.M. Arnold

## Abstract

**Background:** The placenta serves a critical role in nutrient uptake and waste elimination for the developing fetus. The placenta is also responsible for the uptake and/or exchange of xenobiotics, including medications, between the maternal and fetal bloodstreams. An estimated 40-80% of women take medications or drugs during pregnancy for a variety of conditions. Very little is understood about fetal drug and nutrient exposure during pregnancy and how it may change over the course of fetal development.

**Objective:** This study aimed to characterize the abundance of transport proteins in placental tissue, which are important in modulating fetal nutrient and drug exposure, over the duration of pregnancy. Mass spectrometry-based global proteomic analysis revealed trends in the expression of thousands of proteins throughout gestation. Focusing on the membrane-associated proteome enabled an increased emphasis on the solute carrier and ATP-binding cassette families of transporter proteins that are critical for nutrient and xenobiotic transport across the maternal-fetal barrier.

**Study Design:** Using data-independent acquisition proteomics, relative abundance of proteins in placental tissue samples was profiled across all three trimesters of pregnancy (Trimester 1 = 16, Trimester 2 = 9, and Term = 9). Membrane fractions were generated to enrich membrane-associated proteins for proteomic analysis. Placental samples were grouped into randomized batches for membrane fraction generation and mass spectrometry analysis. Proteomic search results from each batch were imported into the R programming environment from Skyline, concatenated, and normalized as one data set for downstream analysis.

**Results:** A total of 6,331 proteins were detected across all samples with 4,210 proteins identified in every sample. Pathway analysis revealed that as gestational age increases, membrane-associated proteins involved in more complex metabolic pathways increase in relative abundance while those involved in extracellular remodeling events and simple organic ion transport tended to decrease. A total of 139 solute carrier and ATP-binding cassette transport proteins were identified in all samples, and 80 were identified in every sample. In general, membrane-associated proteins, including solute carrier and ATP-binding cassette transport proteins, were significantly enriched in placental tissue collected during early gestation compared to term placental tissue.

**Conclusion:** This study presents a comprehensive profiling of membrane-associated proteomic changes during gestation and identifies significant gestational age associated abundance changes at the protein level in several transport protein families. The application of data-independent acquisition global proteomic techniques enabled in-depth analysis of thousands of proteomic changes across pregnancy in a single experiment. These data provide critical information to support future studies into the understanding of fetal exposure to xenobiotics and nutrients circulating in the maternal bloodstream.

## Introduction

The placenta represents the primary conduit between the fetus and mother for the supply of nutrients and metabolic intermediates required for fetal growth. In addition, the placenta functions to remove waste products from the fetal compartment and to reduce fetal exposure to undesirable compounds, including many xenobiotics that may be consumed by the mother. A recent study estimated that ∼73% of pregnant women in the United States take at least one medication (excluding vitamins, supplements and vaccines) during pregnancy with 55% taking a medication during the first trimester (1). Additionally, the use of prescription medications in pregnant women continues to increase despite a lack of relevant maternal-fetal safety, efficacy and pharmacokinetic data for many medications (2). The placenta also serves as a critical site for nutrient transport between the maternal and fetal bloodstreams. During gestation, the placenta undergoes considerable remodeling to accommodate the growth and development of the fetus (3). Changes in nutrient transporter expression are required to ensure proper maturation of the fetus, and deficiencies or imbalances in transporter protein expression have been linked to fetal growth abnormalities such as intrauterine growth restriction or fetal macrosomia (4, 5).

The two primary classes of transport proteins responsible for transport of molecules across the placental barrier are the solute-carrier (SLC) and ATP-binding cassette (ABC) superfamilies of transport proteins (6). Several previous studies have applied RNA-based methods to establish the presence and relative abundance of SLC and ABC family members present in placental tissue (6, 7). While these datasets are informative, there are well established limitations in translating RNA abundance to tissue protein concentrations (8, 9). Targeted proteomics with liquid chromatography tandem mass spectrometry (LC-MS/MS) was previously used to quantify the protein concentrations of several SLC (*n* = 13) and ABC (*n* = 8) transporters across gestation in the placenta (10). While the targeted LC-MS/MS approach was highly selective and sensitive, the approach only generated data for select protein targets incorporated into the method. To increase the number of proteins measured in placental samples, recent studies applied data-dependent acquisition (DDA) proteomics to establish protein profiles for transporters in placental tissue (11–13). While these DDA studies dramatically increased the number of proteins identified in placenta tissue samples compared to targeted studies, the DDA studies only incorporated placental tissue collected at term. In addition, DDA proteomics suffers from inherent limitations in the method such as relying on a “Top *n*” method for selecting precursor peptides, which may introduce a bias toward only the most abundant or most easily ionizable proteins and peptides present in the sample (14). Further, DDA proteomics does not provide relative quantification in complex samples. Given the large number of human SLC and ABC transporters (∼450), and the lack of published protein-level data for a vast majority of these proteins in the placenta, untargeted proteomics with data independent acquisition (DIA) is an ideal approach to identify and quantify transporters. Fragments of all precursor ions present across a chromatographic peak are generated by DIA, leading to increased coverage of lower abundance peptides and proteins (15–17).

The primary goal of this work was to profile the abundance of placental xenobiotic and nutrient transporters over the course of gestation using a DIA proteomics approach. Building a deeper understanding of these proteins at the interface between the fetal and maternal bloodstreams will inform future studies designed to address questions regarding the safety and efficacy of therapeutics and nutrient supplements used during pregnancy.

## Materials and Methods

### Sample collection

The collection of placentae from uncomplicated pregnancies was approved and classified as nonhuman subject research by the Institutional Review Board of the University of Washington. All tissues were collected before 2024. Drug, medication, alcohol, and tobacco use information was not available for all subjects. Ethnicity information was not available for all donors, nor was parity information or whether each sample was from a singleton pregnancy. Fetal sex was only available for a portion of the donors **(Table 1)**. Placentae were divided into three age groups (days ± SD) based on trimester of pregnancy. First trimester (T1: 75 ± 14 days, *n* = 16) and second trimester (T2: 118 ± 13.5 days, *n* = 9) placentae were obtained from University of Washington Birth Defect Research Laboratory. Term (*n* = 9) placentae were obtained from the Labor and Delivery Unit at the University of Washington (18). After collection, tissues were immediately snap frozen and stored at −80°C. Gestational ages for T1 and T2 samples were determined by adding 14 days to the age based on fetal foot length or based on self-reported last menstrual period (19).

**Table 1.** Demographic information for placental tissue donors used in this study. A total of 34 samples were incorporated into the study design spanning all three trimesters.

| **Demographic Category** | *n* |
| --- | --- |
| **Sex** |  |
| Male | 7 |
| Female | 7 |
| Unknown | 20 |
| **Ethnicity/Race** |  |
| African American | 3 |
| Asian | 5 |
| Caucasian/White | 10 |
| Hispanic | 1 |
| Not Available | 15 |
| **Drug Use** |  |
| Reported | 2 |
| Not Available | 32 |

### Isolation of placental membrane fractions

Membrane fractions were generated from placental tissue samples using a standard microsomal preparation method (20). Briefly, frozen tissue from each sample was diced, homogenized on ice, and sonicated in ice cold KPi sucrose buffer (50 mM potassium phosphate pH 7.4, 250 mM sucrose (Fisher, S5-500), 1 mM EDTA (Sigma, E9884), 1x cOmplete Protease Inhibitor Cocktail (Roche, 11697498001)). Homogenates were centrifuged at 10,000 x g at 4°C for 30 minutes. The supernatant was collected and centrifuged at 100,000 x g at 4°C for 1 hour. After ultracentrifugation, the supernatant was removed and membrane fractions were homogenized on ice in resuspension buffer (50 mM potassium phosphate, pH 7.4 and 20% v/v of glycerol (Sigma, G7757)). Resuspended pellets were aliquoted and stored at -80°C until analysis. Additional details are available in Supplemental Materials.

### Proteomic sample preparation

Protein concentration of each resuspended membrane fraction was determined using a Pierce Bicinchoninic Acid assay (Pierce, 23277) and 50 µg of each sample was placed in 36 µl Tris-HCl (pH 7.4, Fisher, BP153-500) for protein digestion. Samples were heat denatured, reduced using dithiothreitol (Pierce, A39255), and alkylated with iodoacetamide (Pierce, A39271) in a final volume of 50 µl. Samples were processed using a modified single-pot, solid-phase-enhanced sample preparation technique (SP3) using a KingFisher Flex (Thermo Fisher Scientific) (21). After trypsin digestion (4 hours at 37°C) and bead elution step, digested peptides were centrifuged at 20,800 x g for 10 minutes at 4°C. Peptides were dried using a vacuum concentrator (Savant SpeedVac SPD 1030, Thermo Fisher) and stored at -80°C until proteomic analysis. Additional details about the sample preparation method are available in the Supplemental Materials.

### Data independent acquisition mass spectrometry and data processing

Samples were randomized into three batches for proteomic sample preparation and data independent acquisition (DIA) analysis following a published batching scheme (22). A standard library-free gas-phase fractionation (GPF) DIA acquisition and analysis workflow was used to analyze the samples in each batch (**Supplemental Table 2**) (23, 24). Raw files were converted into the mzML format using ProteoWizard (v. 3.0.23318) and processed using EncyclopeDIA (24). The whole human proteome FASTA file (Uniprot, downloaded 11/2023) was used for proteomic searches as well as the background proteome file in Skyline with trypsin set as the protease allowing for one missed cleavage (25). Peptide matches to proteins were filtered in Skyline and required at least one unique peptide/protein with at least three transitions/precursor and a minimum dotp score of 0.8. Additional details are available in Supplemental Materials.

### Proteomic data analysis

Using a predefined Skyline export template, filtered Skyline results for each batch were imported and processed using the MS Stats pipeline running in R Studio (v 2024.12.1, running R v 4.5.3) · (26). Individual batch files were merged after import prior to processing. Tukey’s Mean Polish was applied to summarize peptide level data which was then quantile normalized and log-transformed into relative abundance values for each protein identified. The limma statistical analysis package was used for batch correction and differential analysis of protein level data · (27). Volcano plots were generated using R. Gene ontology (GO) analysis was performed using ShinyGO (ver. 0.85) and incorporated proteins identified as significantly, differentially expressed in each trimester (adjusted *p*-value < 0.05 and log2 fold change cutoff < -1 or >1) (28). Principal component analysis (PCA) using PCA Tools was used to perform unsupervised analysis (29). Additional details are available in Supplemental Materials.

### Statistical testing

Empirical Bayes moderation (eBayes) was used to compute moderated t-statistics and corresponding raw *p*-values for each trimester comparison. Raw *p*-values were then adjusted for multiple testing using limma’s default Benjamini–Hochberg false discovery rate procedure. Protein mean abundance plots were generated in R using the ggplot2 package. For each protein, the previously computed abundances were grouped by trimester and the statistical significance for each trimester comparison was taken from the adjusted *p*-values applied during the limma differential analysis workflow.

### Data records

The complete datasets for the DIA studies are available via Panorama Public (https://panoramaweb.org/uwtec_placenta_transporterDIA.url) and were assigned the ProteomeXchange ID: PXD080148(30, 31). Experimental data are provided at multiple levels of processing, from raw MS data to normalized peptide-and protein-level quantities suitable for downstream analysis

## Results

### Relative abundance changes and pathway analysis during fetal development for all observed placental proteins

Membrane fractions generated from 34 placental samples were analyzed using library-free DIA-MS proteomics. No significant difference in microsomal protein yield was observed based on gestational age of the samples. Across all samples, 6,331 proteins passed all filtering criteria with 4,210 proteins measured in every sample. The mean age (± SD) of T1 samples was 75 ± 14 days, T2 samples had a mean age of 118 ± 13.5 days, and all Term samples were considered full term at 287 days. **Figure 1** shows differential analysis of protein abundance by trimester. Even with the relatively small age difference between the T1 and T2 samples, more than fifty proteins were significantly differentially (2-fold difference) expressed between T1 and T2 (**Figure 1A**). The largest number of significantly differentially expressed membrane-associated proteins was identified when comparing T1 and Term samples (**Figure 1C-D**) which also represents the largest age gap between samples. The full list of proteins with associated fold changes and *p*-values by trimester is available in **Supplemental Table 3**.

**Figure 1.**
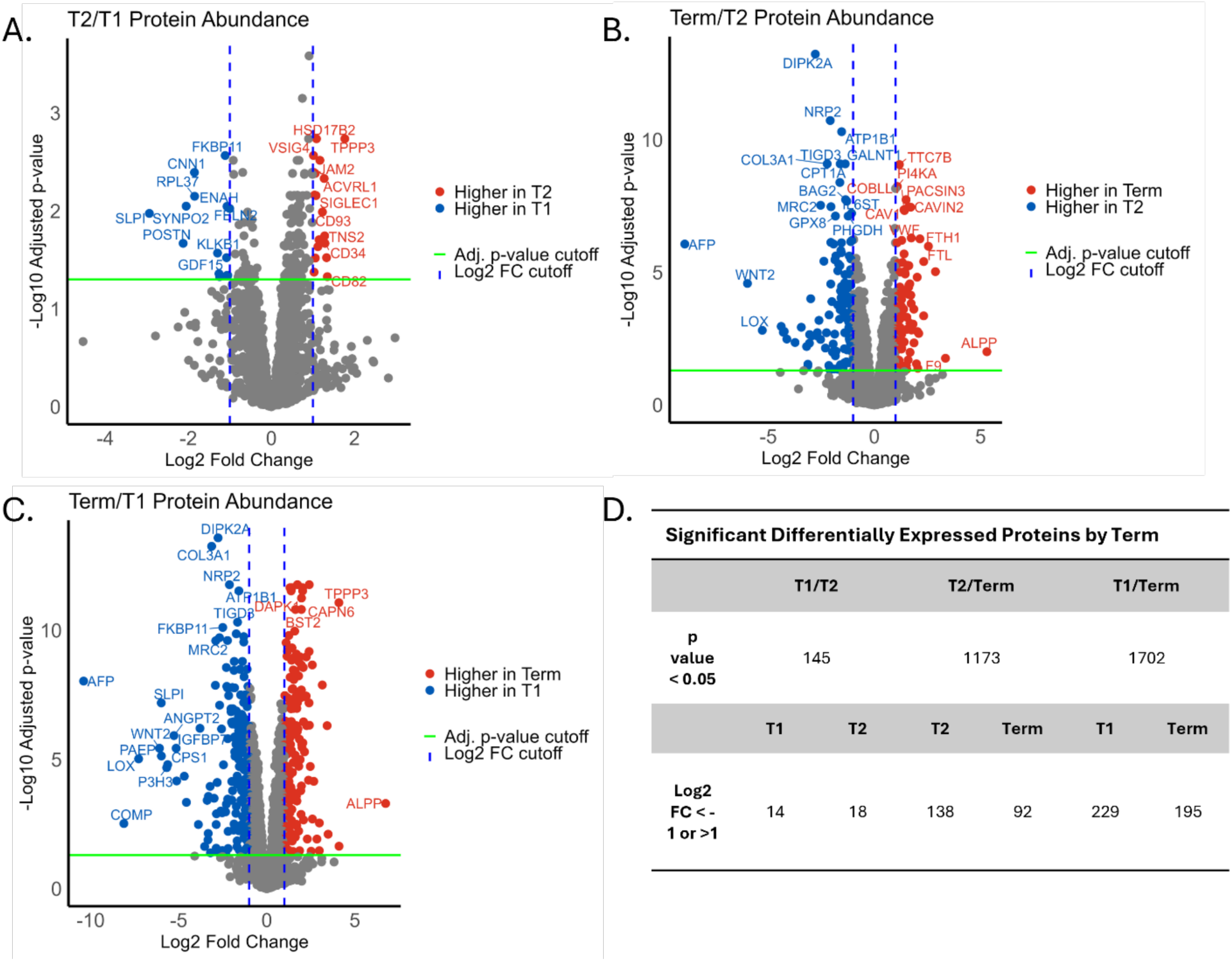
Volcano plots for all microsomal proteins passing the search filter criteria between trimesters (T1 *n* = 16, T2 *n* = 9, Term *n* = 9). The three comparisons for differential analysis included T2/T1 (A), Term/T2 (B), and Term/T1 (C). Table (D) displays the numerical distribution of proteins that pass statistical criteria (adjusted *p*-value < 0.05) and log fold change cutoff (Log2FC < -1 or >1) cutoff for each trimester. Adjusted *p*-values were determined using the Benjamini-Hochberg procedure on the raw *p*-values determined by differential analysis.

GO analysis was used to identify the top ten biological processes for each trimester associated with the differentially expressed proteins (**Figure 2A-C**). During early development, upregulated biological processes consist largely of extracellular reorganization and matrix assembly and preparative functions for the growth of the placenta throughout pregnancy such as mitochondrial translation and blood vessel and vasculature development. In the second trimester, as the placenta undergoes reorganization, more complex processes such as lipid and steroid metabolism, cholesterol metabolic processes, and tube development for increasing vascularization become prominent. Finally, at term, complex networks such as the regulation of cell motility and migration, circulatory system development, and cytoskeleton organization processes were enriched. Vasculature, blood vessel, and circulatory system development pathways also continue to be prominently upregulated.

**Figure 2.**
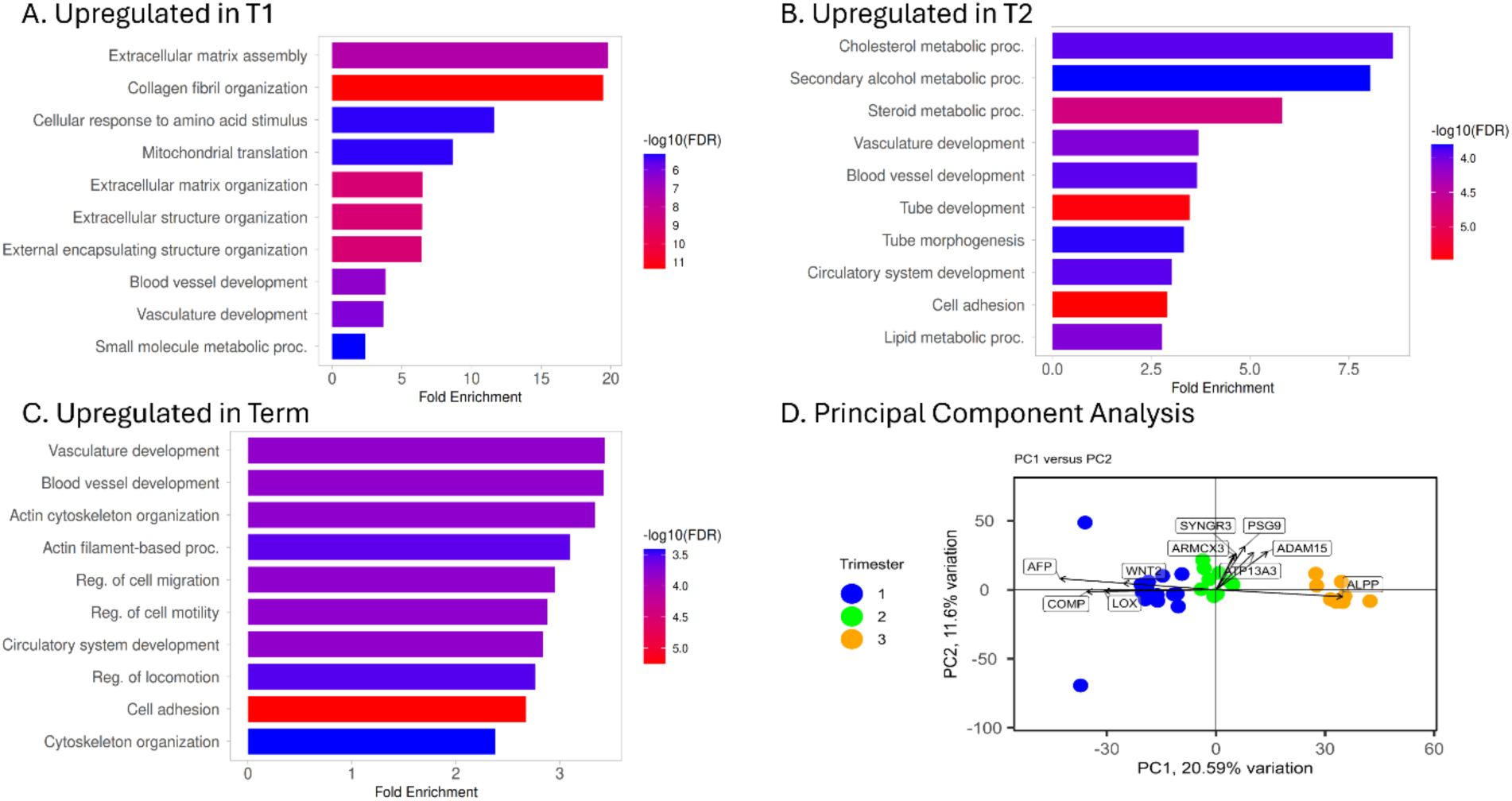
Significant differentially regulated biological processes associated with each trimester. Membrane-associated proteins identified as significantly altered in T1 (A), T2 (B), and Term (C) were analyzed with ShinyGO using a 0.05 false discovery rate. Panel D shows the first two principal components (PCs) from principal component analysis of the proteins detected across every sample in the data set. Arrows indicate each labeled protein’s contribution to the principal components, with arrow direction showing association with each PC and arrow length reflecting contribution magnitude.

PCA was performed with proteins identified in every microsomal sample, and a biplot for the first two principal components (PC) is shown in **Figure 2D** (29). The samples group by gestational age, as denoted by the color coding for each trimester, reflecting the previously acknowledged small age gap between T1 and T2 samples and the larger age difference between the Term samples and T1 and T2. Arrows indicate the PC loadings, representing the contribution of each labeled protein to PC1 and PC2. As expected, PLAP (Alkaline phosphatase, placental) and AFP (Alpha-fetoprotein) differentiate strongly between early and late gestational ages (**Figure 2D and Supplemental Figure 1**). There were relatively subtle changes in the abundance of PSG9 (Pregnancy-specific beta-1-glycoprotein 9) and ADAM15 (Disintegrin and metalloproteinase domain-containing protein 15) as the placenta matures.

### Transporter specific changes highlight metabolic trends present in each trimester

Differential analysis plots with only SLC and ABC family member transport proteins are shown in **Figure 3**. As previously observed at the whole membrane-associated proteome level, the largest abundance differences exist between T1 and Term samples, with more transporters identified as having significantly higher abundance in T1 (**Figure 3C**). Most transport proteins changing over gestation are in the SLC family, with only ABCB1 (P-gp) and ABCG2 (BCRP) from the ABC family significantly higher in T1 and T2 over Term (**Figure 3B and 3C)**.

**Figure 3.**
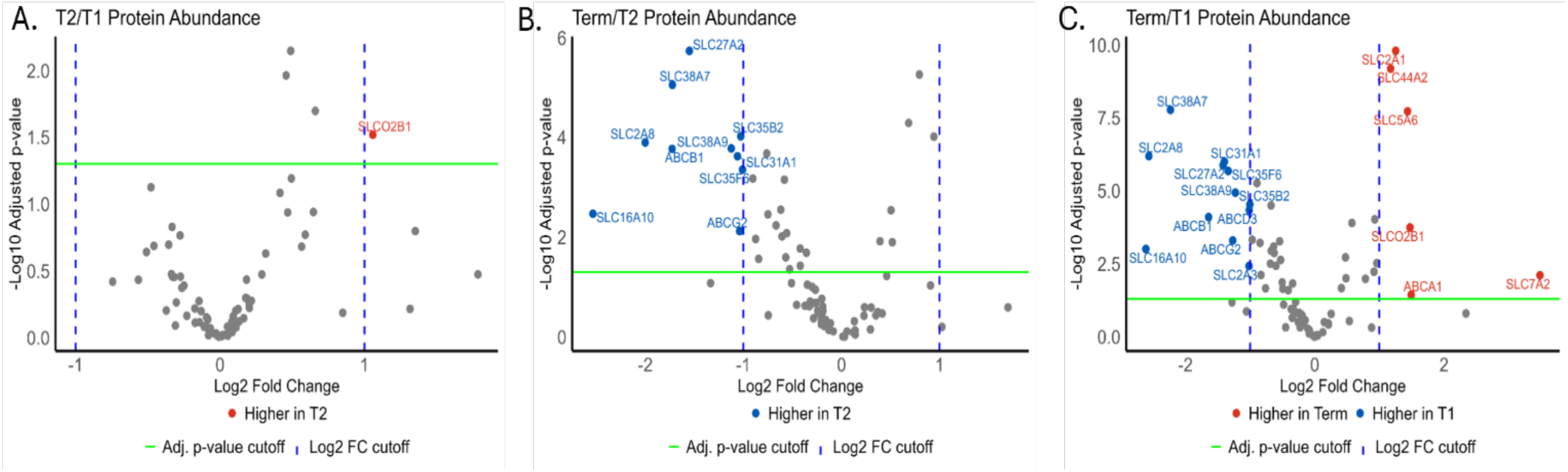
Volcano plots for the SLC and ABC families of transport proteins. The three comparisons for differential analysis included T2/T1 (A), Term/T2 (B), and Term/T1 (C) (T1 *n* = 16, T2 *n* = 9, Term *n* = 9). Proteins that passed statistical criteria (adjusted *p*-value < 0.05) and log fold change cutoff (Log2FC < -1 or >1) for each trimester (green line) are displayed as blue or red points. Adjusted *p*-values were determined using the Benjamini-Hochberg procedure on the raw *p*-values determined by differential analysis of the entire data set.

A more granular examination of individual SLC proteins based on the normalized abundance values per trimester is shown in **Figure 4**. In **Figure 4A**, two SLCs involved in organic anion and vitamin transport, SLCO2A1 (OATPA1) and SLC5A6 (SMVT) respectively, increase in abundance from T1 to Term, while SLC27A1 (FATP1) and 27A2 (FATP2) highlight increased fatty acid transport during the second trimester. **Figure 4B** shows changes in the SLC2 family of glucose transporters throughout fetal development. **Figure 4C** highlights variation in abundance for specific amino acid transporters in placental tissue. In general, the observed ABC family transporters (**Figure 5**) exhibit fewer changes throughout gestation and no large abundance differences akin to those observed in the SLC families. **Supplemental Table 3** contains a full list of SLC and ABC transporters identified in each trimester.

**Figure 4.**
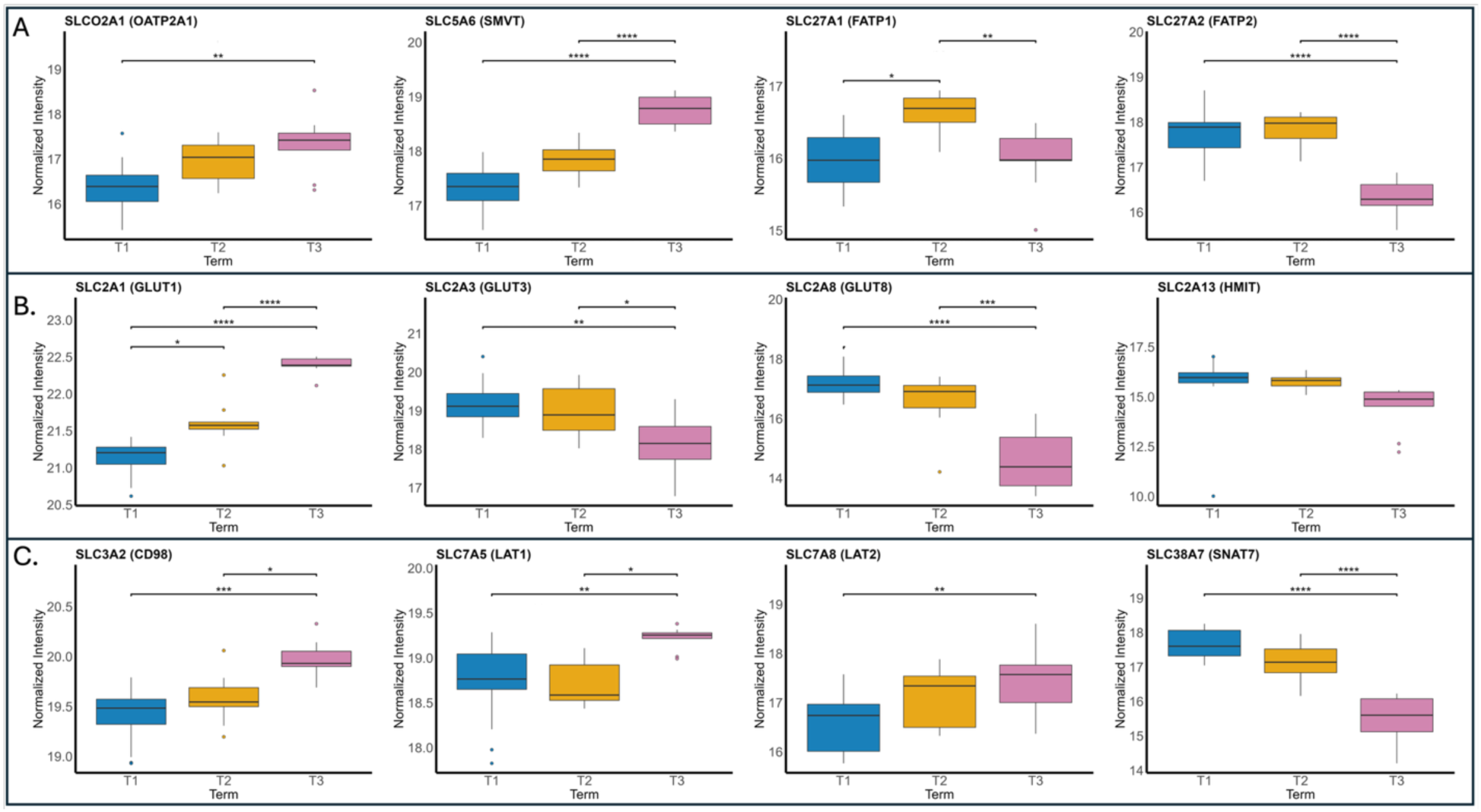
Individual protein plots comparing mean abundance values of SLC transporter proteins across trimesters. Individual proteins of interest are plotted to illustrate significant trends for known pathways and/or classes of transporters previously established to be important or significantly changing across trimesters (T1 *n* = 16, T2 *n* = 9, Term *n* = 9). Mean abundance values for members of the vitamin and fatty acid transport families (A), glucose transporter family (B) and amino acid transporter changes (C) are shown grouped by trimester. Black bars represent the median value of the normalized, log-transformed, batch-corrected abundance value per trimester. Boxes denote the interquartile range between 25% and 75% of the normalized intensity range. Whiskers extend to cover 1.5x the interquartile range and dots represent outlier samples. Statistical significance was computed using the Benjamini-Hochberg method applied to the raw *p*-values calculated from empirical Bayes analysis after linear modeling trends in the data using trimester comparisons as contrasts. * = *p* < 0.05, ** = *p* <0.01, *** = *p* < 0.001, and **** = *p* < 0.0001.

**Figure 5.**
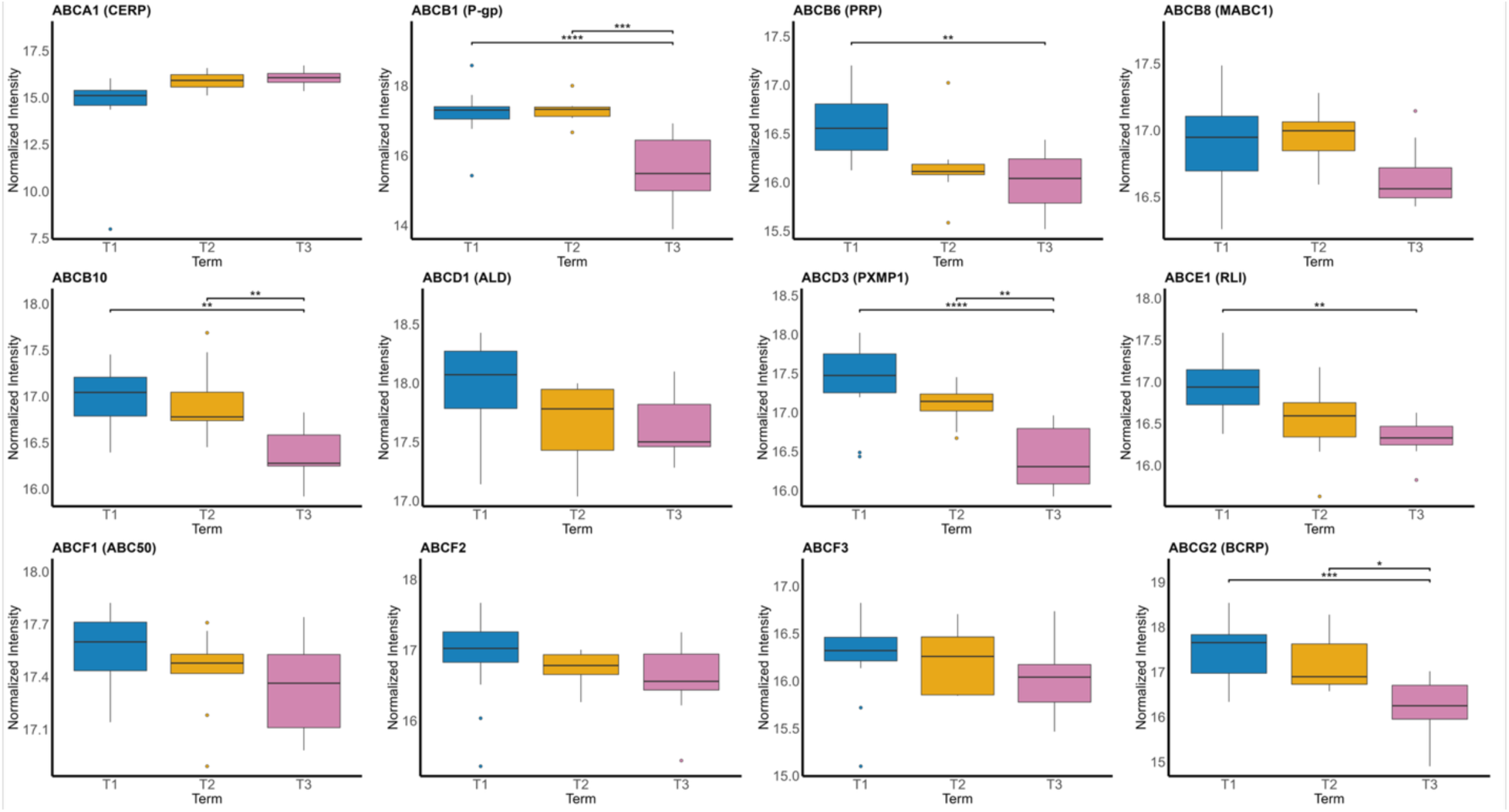
Individual protein plots comparing mean values of ABC transporter proteins across trimesters. All the proteins from the ABC family that were found in every sample are shown. Black bars represent the median value of the normalized, log-transformed, batch-corrected abundance value per trimester (T1 *n* = 16, T2 *n* = 9, Term *n* = 9). Boxes denote the interquartile range between 25% and 75% of the normalized intensity range. Whiskers extend to cover 1.5x the interquartile range and dots represent outlier samples. Statistical significance was computed using the Benjamini-Hochberg method applied to the raw *p*-values calculated from empirical Bayes analysis after linear modeling trends in the data using trimester comparisons as contrasts. * = *p* < 0.05, ** = *p* <0.01, *** = *p* < 0.001, and **** = *p* < 0.0001.

### Comment

To the best of our knowledge, this study represents the first global proteomic characterization of placental membrane-associated proteins using DIA-MS proteomics. The overall goal of this work was to examine protein abundance changes across gestation with a special focus on SLC and ABC transport proteins. Of the 6,331 proteins identified, 139 were from either the SLC or ABC families. Since the DIA data were collected in separate batches, the focus was narrowed to the 4,210 proteins measured in every sample across all three batches which yielded a total of 80 transport proteins, 17 of which (14 SLC and 3 ABC) were identified as significantly altered between trimesters during fetal development (**Figure 3**).

Differential analysis with all observed placental membrane proteins revealed that the main characteristic associated with changes in placental protein abundance is the gestational age **(Figure 1)**. GO analysis indicated biological processes associated with each trimester correspond to increasingly complex tissue growth processes and metabolic needs, reflecting the changing metabolic demands of the growing fetus and overall structural changes to the placenta itself as it grows **(Figure 2**). PCA highlighted specific proteins, such as AFP and PLAP, which are known to display opposing expression profiles during gestation, illustrating the relationship of each protein to either end of the sample age distribution **(Figure 2D)** (32, 33).

SLC transporters enriched in Term samples compared to T1 and T2 generally encompass the movement of more complex molecules across the cell membrane, such as those involved in vitamin transport. For example, SLC5A6, which is responsible for pantothenate, biotin, lipoate and iodide transport, and SLCO2A1, responsible for organic anion transport, are enriched in Term samples compared to T1 (**Figure 4A)**. SLC27A1 and SLC27A2 protein abundance were enriched in T2 samples which align with increased fatty acid uptake by the fetus during this stage of development and increased fatty acid deposition in trophoblasts prior to the onset of high fetal fatty acid demand (**Figure 4A**) (34, 35). Together, these observed patterns align with the increased complexity of energy and metabolic demands from the fetus over the course of development.

Along with vitamin and fatty acid transport, SLC proteins involved in carbohydrate transport displayed distinct changes in relative abundance across fetal development. Glucose serves as the primary energy source for the fetus, and the energy requirements for fetal development increase over the course of gestation. The fetus has a limited capacity to synthesize glucose, and the enzymes in the gluconeogenic pathway display markedly decreased activity compared with post-natal levels (36, 37). Fetal glucose is predominantly acquired from the mother, and placental SLC2A1 (GLUT1) expression has been shown to increase ∼2-fold between 16-22 weeks and 27-30 weeks of gestation (38, 39). Similar to previous reports, the mean abundance of SLC2A1 in our proteomics study dramatically increased over the course of gestation which reflects the increasing demand for glucose to support fetal metabolism and growth (**Figure 4B)**.

In contrast to SLC2A1, SLC2A3 (GLUT3) abundance in our proteomics study was observed to trend downwards through fetal development, and this aligns with previously published western blot data (40). SLC2A3 upregulation has been previously associated with hypoxic conditions, such as those known to exist during the first trimester while the placenta is poorly vascularized and oxygenated (41). SLC2A3 is a rapid, high-volume glucose transporter with high affinity for glucose, which would promote placental glucose uptake under adverse conditions (40, 42). As the placenta grows concomitant with fetal development and becomes more vascularized, SLC2A3 expression driven by hypoxic conditions decreases and is likely compensated by increase in SLC2A1 abundance as glucose needs increase as the placenta develops.

SLC2A8 (GLUT8) has been shown to localize primarily to intracellular membranes in multiple trophoblast cell types in the placenta (43). The role of SLC2A8 in fetal development remains largely unknown, with a recently published RNA interference study suggesting mostly indirect effects on human trophoblast glucose uptake (44). The protein-level data presented here suggests a gestational-age-associated downregulation of SLC2A8 throughout fetal development and further studies into the cause and effect of this change are warranted. Finally, in the glucose transporter family, the expression of SLC2A13 (HMIT) exhibited no significant changes over gestation. SLC2A13 expression has been previously confirmed in placental tissue and studied in the context of gestational diabetes mellitus onset leading to the downregulation of inositol transporters, including SLC2A13, in response to elevated maternal glucose levels (45).

As observed with glucose transporters, the present study identified gestational-age-associated changes in amino acid transporters (**Figure 4C**). SLC38A7 (SNAT7), a sodium-coupled neutral amino acid transporter involved in lysosomal amino acid transport and mTORC1 activation, was observed to have higher expression in T1 compared to Term samples. SLC3A2 (CD98) can form a dimer with either SLC7A5 (LAT1) or SLC7A8 (LAT2) for amino acid transport in the placenta (46). Together these complexes are focused on transporting large, neutral amino acids. The mTORC1 pathway has been identified as a critical link for sensing amino acid availability in placental trophoblasts and is therefore important in regulating the maternal supply of amino acids for fetal growth (47).

Several ABC family members exhibited gestational-age-related changes, with most demonstrating a downward trend from T1 to Term **(Figure 5)** . When comparing membrane fractions generated from T1 and Term placentas, a previous targeted proteomics study reported a 55% and 69% reduction in ABCG2 and ABCB1 expression, respectively (10). These two transporters were also quantified in our global proteomics study, and we identified a nearly identical average decline in abundance from first trimester to term for ABCG2 (59%) and ABCB1 (68%) (**Figure 5)**. A third apical membrane-associated protein, SLC6A4 (SERT) was also detected in both the targeted study and the current global proteomics dataset, again with a relatively modest decline in expression from first trimester to term (data not shown). The similar trends across gestation observed with both targeted proteomics (MRM) and global proteomics (DIA) methods further support the quantitative protein level data generated with our DIA-only approach.

This study lays the groundwork for further advancements in the understanding of the membrane-associated proteome during fetal development. Specifically, by highlighting relative abundance changes in several subfamilies of SLC and ABC proteins throughout gestation, the proteomic dataset provides an improved understanding of how changing metabolic demands associated with gestational development influence placental transporter levels. Furthermore, these data will inform future studies on potential nutrient and xenobiotic transport across the placental barrier and improve the understanding of the potential implications on fetal exposure to exogenous compounds consumed by the mother during pregnancy. Future studies to model and simulate the maternal-fetal distribution of transporter substrates will require the abundance of transporters in the entire placenta (49–51). We used the PLAP value for each sample to reflect placental size and estimated the total relative abundance of select SLC transporters during fetal development (**Supplemental Figure 3**). Overall, there is a consistent gestational age–dependent increase in abundance for all transporters. This observation was expected due to the dramatic increase in placental size with gestation compared to modest changes in relative protein abundance.

A major strength of this study is the quantitative data generated with DIA proteomics for thousands of analytes in a single experiment which yields a rich dataset that can provide information on proteins beyond the specific targets of the current study. Proteomic data presented in the current work are available on Panorama Public (https://panoramaweb.org/uwtec_placenta_transporterDIA.url) and were assigned the ProteomeXchange ID: PXD080148 (30, 31). These resources allow anyone to conduct their own analysis with their targets of interest. In the present study, multiple members of each transporter type were interrogated in a single experiment without the need for complex labeling schemes or multiplexing limitations that would arise from antibody-based methods. A limitation to this study includes lack of an appropriate number of samples with a confirmed sex, which prevented evaluation of the contribution of fetal sex to the membrane-associated placental proteome in this study. Additional study metadata were missing that would have enriched this analysis such as donor age, parity, singleton status of each pregnancy, and tobacco or other medication or substance use information.

In conclusion, this work provides insight into specific transport proteins and their regulation throughout pregnancy. While many of the primary changes in transporter abundance in the placenta across gestation are associated with fetal growth and metabolism, there are subtle patterns within families of transport proteins that do not display one single trend.

## Acknowledgements

We thank Dr. Drake Russell for his assistance developing the protocol to generate placental membrane fractions.

## Author Contributions

**Eric Weaver**: Formal analysis, Investigation, Visualization, Writing - Original Draft **Ariel Topletz-Erickson**: Resources **Nina Isoherranen**: Resources, Writing - Review & Editing, **Jashvant Unadkat**: Conceptualization, Resources, Writing - Review & Editing, Funding acquisition **Samuel Arnold**: Conceptualization, Resources, Writing - Review & Editing, Supervision, Project administration, Funding acquisition

