## Supplemental Material for "Proteomic profiling of xenobiotic and nutrient transporters in human placenta of different gestational ages"

### Materials and Methods

#### Isolation of placental membrane fractions

Membrane fractions were generated from 34 placental tissue samples using a standard microsome preparation method (1). Between 1.5 and 3.5 grams of frozen tissue from each sample was diced into ~1 $mm^3$ portions in a weigh boat set on a pre-chilled aluminum block on a bed of dry ice. Diced tissues were placed in a 50 mL conical tube (Falcon, 352070), resuspended in approximately 30 mLs of ice cold 1x phosphate buffered saline (PBS) (Gibco, 10010023) and rinsed by gently rocking the tube back and forth several times. The tissue pieces were collected into a 100 µm cell strainer (Falcon, 352360) rinsed with 3x 10 mL washes of ice cold 1x PBS. Further washes with PBS were necessary for tissues that contained higher amounts of visible blood. Samples were transferred to a 10 mL Dounce homogenizer (douncer) and resuspended in 3.5 mLs per 1 g of starting tissue weight of ice cold KPi Sucrose buffer (50 mM potassium phosphate pH 7.4, 250 mM sucrose (Fisher, S5-500), 1 mM EDTA (Sigma, E9884), 1x cOmplete Protease Inhibitor Cocktail (Roche, 11697498001)). Tissues were homogenized on ice with a drill powered pestle that was passed through the tissue in the douncer 4-6 times, or until no solid pieces of tissue remained. After homogenization, a probe sonicator (Fisher Scientific) set to 80% power was passed through the homogenized tissue four times, 20 seconds each, with a one-minute rest in between passes. The homogenate remained on ice for the duration of the sonication. Sonicated homogenate was centrifuged at 10,000 x g at 4°C for 30 minutes. The supernatant was collected and centrifuged at 100,000 x g at 4°C for 1 hour. After ultracentrifugation, supernatant was removed and the pellet was rinsed once with fresh, ice cold KPi glycerol buffer (50 mM potassium phosphate, pH 7.4 and 20% v/v of glycerol (Sigma, G7757)). A plastic spatula was used to transfer the pellet from the centrifuge tube to a clean, pre-chilled douncer. Once transferred, 1 mL per 1 g of starting tissue weight of KPi glycerol was added to the douncer and the pellets were homogenized by hand with a pestle until the pellet was completely resuspended in the buffer. Resuspended pellets were aliquoted and stored at -80°C until analysis. All homogenization steps were carried out on ice.

**Proteomic Sample Preparation**

Samples were prepared in three randomly selected batches consisting of at least one sample from each gestational age (Supplemental Table 2). Membrane fraction pellets were processed using a modified single-pot, solid-phase-enhanced sample preparation technique (SP3) using a KingFisher Flex (Thermo Fisher Scientific) (2). Briefly, each sample was thawed on ice and total microsomal protein concentration was determined using a BCA assay (Pierce, 23277). Samples were normalized to a starting protein concentration of 50 µg in a total volume of 36 µL. A total of 800 ng of yeast enolase was added to each sample as an internal digest control. Samples were denatured by heating to 95°C in a thermomixer (Eppendorf) with gentle mixing for 5 minutes, then 100 mM dithiothreital (DTT, Pierce, A39255) was added to a final concentration of 20 mM, vortexed, and then incubated at room temperature for 20 minutes. After denaturation and reduction, the samples were alkylated with 40 mM iodacetamide (IAA, Pierce, A39271) in the dark at room temperature for 15 minutes. MagReSyn Hydroxyl beads (Resyn Biosciences, MR-HYX010) at their stock concentration (20 µg/mL) were added to a deep-well 96 well Kingfisher plate at a ratio of 250 µg of beads to 50 µg of protein. After alkylation, samples were added to the corresponding wells with beads and gently mixed by pipetting up and down several times. A volume of 100% acetonitrile (ACN) was added to each sample well to reach a 70% final concentration and allowed to precipitate at room temperature for 10 minutes.

After sample mixing and binding with the hydroxyl magnetic beads, the complexed beads and microsomal protein were loaded onto the KingFisher Flex and washed in 95% ACN three times and 70% ethanol twice. A final digest/elution plate was prepared using Tris-HCl (pH 7.4) and trypsin (Pierce, 90058) to make a 20:1 working ratio of protein:enzyme in a final volume of 50 µL per well. Protein and beads were then eluted into the Tris/trypsin mixture in the elution plate which was subsequently heated to 37°C and digested for 4 hours with intermittent mixing. After the digestion and elution step, the magnetic beads were removed. Digested peptides were moved from the Kingfisher plate into individual Eppendorf tubes and spun at 20,800 x g for 10 minutes at 4°C to pellet any residual beads left in the samples. After spinning, samples were transferred to fresh tubes and evaporated to dryness using a vacuum concentrator (Savant SpeedVac SPD 1030, Thermo Fisher). Dried peptides were stored at -80°C until proteomic analysis.

**Data independent acquisition mass spectrometry**

Sample injection order was randomized within each batch for DIA analysis following a previously published batching scheme (3). Dried samples for each batch were placed on ice and resuspended in a solution of 2% ACN/0.1% formic acid at a final concentration of 1 µg/µL. Individual samples were mixed with Pierce Peptide Retention Time Calibration (PRTC, Pierce, 88321) mixture at a final concentration of 100 fmol/µL. Gas phase-fractionated library (GPF library) samples were generated by combining equal amounts of each sample in a batch and mixing with PRTC. All samples representing each trimester in each batch (T1, T2, Term) were also pooled separately and mixed with PRTC to serve as intrabatch control samples to monitor data quality and MS performance within each batch. An additional interbatch control sample consisting of a cytosolic fraction of the membrane fraction isolation high-speed centrifugation was also added to each batch and included in the GPF library pool to monitor data quality and MS performance across batches (3).

Proteomic analysis using DIA was carried out on an Orbitrap Exploris 480 (Thermo Fisher Scientific) coupled to a Vanquish Neo UHPLC (Thermo Fisher Scientific). A Bruker PepSep 30 cm x 150 µm column (Bruker, 1895833) with a 5 mm x 300 µm guard column (Thermo Scientific, 174500) was used to separate peptides. One microgram of sample was injected and analyzed using a trap-and-elute method, peptides were separated using a linear gradient of 4% solvent B (80% ACN with 0.1% formic acid) to 50% solvent B over 42 minutes. A previously published, library-free DIA acquisition scheme was used to analyze each batch (4, 5). Briefly, GPF chromatogram libraries were acquired using the pooled sample in each batch analyzed by six sequential injections in 100 *m/z* windows ranging from 395 to 1005 *m/z*. Each injection used a "narrow window" 4 *m/z*-wide DIA spectra at 30k resolution, an automatic gain control (AGC) target of 1 x $10^6$ with the maximum inject time set to "Auto" and a normalized collision energy of 27. Two precursor ion spectra were acquired after every 25 DIA spectra. The wide window precursor spectrum spanned 400-1600 *m/z* and narrow window precursor spectrum matching the mass window for each method (i.e., 395-505, 495-605, 595-705, 695-805, 795-905, and 895-1005 *m/z*) were acquired with 60k resolution and a 1 x $10^6$ AGC target using "Auto" to determine AGC fill time. DIA analysis for individual samples was carried out over the 400-1004 *m/z* mass range using "wide window" 8 *m/z*-wide DIA spectra at 30k resolution with an AGC target of 1 x $10^6$, maximum injection time set to "Auto" and a normalized collision energy of 27. Every 25 DIA spectra, a precursor ion spectrum was acquired over the 400-1600 *m/z* mass range at 60k resolution, 1 x $10^6$ AGC target and maximum injection time set to "Auto."

**Data Processing**

DIA data analysis and processing was performed following a previously published workflow (4). Briefly, raw data files were demultiplexed with 10 ppm accuracy and converted to the mzML format using ProteoWizard (v. 3.0.23318) (6). Prosit was used to generate an *in silico* peptide library for the whole human proteome (Uniprot, KB 9966, downloaded 11/2023) with the charge range set at 2-3, one missed cleavage, default *m/z* range, an NCE 27 and a default charge state of three (5, 7). mzML files generated from the GPF library samples were searched against the Prosit-generated spectral database in EncyclopeDIA (version 2.12.30) using the default settings. Search results were filtered using a 1% false discovery rate (FDR) cutoff at the peptide level using Percolator v. 3.01. Peptide search results were saved in the elib file format as a "Chromatogram Library". Next, single injection mzML files were searched against the GPF chromatogram elib using the whole human proteome FASTA file as a background (Uniprot, downloaded 11/2023), search results were filtered using the same criteria used for the chromatogram library searches. Individual sample searches for each batch were combined and saved as an elib file using the "Quant Report" feature. After the initial peptide searches were completed in EncyclopeDIA, Skyline (24.1.0.199) was used to further process and filter the data. Using previously published import settings, the elib quant report file was loaded into Skyline as a spectral library and used to search the demultiplexed mzML files generated from the raw data files for each individual sample injection (4). The whole human proteome FASTA file was used as the background proteome file in Skyline with trypsin set as the protease allowing for 1 missed cleavage. Peptide matches to proteins were filtered in Skyline and required at least one unique peptide/protein with a minimum of 3 transitions/precursor and a minimum dotp score of 0.8. Repeat and duplicate peptides shared between proteins were removed. Peptide uniqueness was enforced at the protein level.

**Proteomic Data Analysis**

Using a predefined Skyline export template, filtered Skyline results for each batch were imported and processed using the MS Stats pipeline running in R Studio (v 2024.12.1, running R v 4.5.3) (8). Individual batch files were merged after import prior to processing with MS Stats. Further processing of the dataset in MS Stats used logarithmic-base 2 transformation of the measured peptide abundances, quantile normalization, and Tukey's Mean Polish summarization to summarize detected features for each protein into a single value representing each protein's measured abundance in the data set.

The summarized, normalized and log-transformed values were then processed using the limma statistical analysis package for batch correction and differential analysis (9). Linear models were fit for each protein and contrasts were set to compare between trimesters using a design matrix and no intercept. Moderated *t*-statistics were computed using empirical Bayes shrinkage with the "treat" function in limma. Raw *p*-values were adjusted for multiple comparisons using the Benjamini-Hochberg procedure. Proteins with an adjusted $p$-value $< 0.05$ and $\log_2$ fold change cutoff $< -1$ or $>1$ were considered significant. Volcano plots were generated using a custom script in R. Gene ontology (GO) analysis was performed on the lists of proteins generated from the differential analysis results using ShinyGO (ver. 0.85) to compare significant proteins from each trimester involved in different biological processes across gestation (10). The top ten pathways with a minimum of five genes at an FDR of 0.05 were set for GO enrichment searches. Principal component analysis (PCA) using PCA Tools was used to perform unsupervised analysis of proteins changing with gestational age (11). Further details regarding data analysis, including R scripts used to process data can be found in Appendix A of Supplemental Information. A table listing all proteins and their significance and fold change per trimester is found in **Supplemental Table 3**.

**Statistical Testing**

Protein mean abundance plots were generated in R using the ggplot2 package. Briefly, the normalized, log-transformed, batch-corrected intensity values for each protein were grouped by trimester. The statistical significance for each comparison was taken from the adjusted $p$-values applied during the limma differential analysis workflow. Comparisons with an adjusted $p < 0.05$ were considered significant. Data are presented as the mean ± SEM.

**Supplemental Discussion**

**Housekeeping and Protein Normalization Discussion**

**Supplemental Figure 1A** shows the mean abundances by trimester for several housekeeping proteins involved in cellular processes that are not expected to vary under the physiological conditions in the present study. As seen in the plots, these proteins are not found to vary significantly by trimester. **Supplemental Figure 1B** shows the normalized relative abundance by trimester of PLAP (Alkaline Phosphatase, Placental) and AFP (Alpha-fetoprotein), known fetal growth markers, with PLAP showing a significant increase from T1 to Term and AFP showing the opposite trend, as expected (12, 13). The plots of these two proteins demonstrate the ability of DIA proteomics to accurately track protein expression associated with placental growth from the first trimester to term.

**Addressing Sample Batched Data Acquisition in DIA Proteomic Analysis**

Differences in SLC and ABC expression patterns between trimesters are largely a reflection of the changing metabolic demands in placental tissue throughout development. The mean gestational ages of the T1 and T2 samples are relatively close together (75 days for T1 and 118 days for T2) which explains the low number of significant differentially expressed proteins in the direct comparison between those groups. When both T1 and T2 are compared separately to Term samples, the larger difference in age captured in each comparisons identifies a larger number of significant differentially expressed proteins between the early gestational age samples and term. Because this analysis was done in separate batches, we elected to focus only on the proteins found in every sample. However, as noted in **Supplemental Table 2**, the total number of proteins and transporters detected across all batches after proteomic filtering is larger than the number of proteins chosen for the final analysis (only proteins detected in all samples across all batches). Because this study is focused on transport proteins, the discrepancy in transport proteins detected in each batch was explored further and shown in **Supplemental Figure 3** using an "Upset Plot" to show overlap and distinctions between the transport proteins detected in each batch. This plot shows the number of transport proteins found in each batch individually and where proteins detected in each individual batch overlap. Overall, batch 3 contained the highest number of transport proteins identified with 125, then batch 2 with 98, and batch 1 with 97. The plot also demonstrates that each batch contains transporter proteins unique to that batch. In the context of DIA proteomics using GPF on-column libraries, this demonstrates a weakness in running a batched analysis. While certain proteins may be detected as unique to one batch, their exclusion from a different batch does not mean that the transporters should be considered "not present" in that batch, rather they should be noted as "not detected." These proteins are likely present in other batches but have no observed unique peptides which exceed detection and filtering criteria.

While 80 transporters were analyzed in-depth because they were detected in all samples (and in all three GPF libraries), the number of transporters found in batch 3 indicates that up to 125 of these proteins could be present, but not detected, in samples from the other batches. While it is computationally possible to search wide-window injections from one batch against an on-column library from a different batch, this should only be done in the context of investigating batch-to-batch variations, as the search results would not be considered valid for reporting results. Future experimental plans could be adapted to avoid running batched analysis, or a searchable library could be established using replicate injections following a data-dependent acquisition (DDA) strategy, which would allow all subsequent single injections to be searched against the same library. However, DDA-based libraries suffer from well-established limitations (4).

**Investigating the Effects of Placental Growth on Protein Levels**

The placenta is a dynamic organ that exhibits rapid growth during gestation. Because of this, placental samples taken at term are sampled from a much larger tissue mass than those sampled from T1 and T2 (14). This size discrepancy presents challenges for comparing protein abundances between trimesters, as placental growth could mask actual trends in protein concentration as a function of overall placental mass. Anoshchenko et al. present a normalization strategy to address this issue that factors in the overall volume of the placenta at different stages of fetal development (15). Data sets acquired by MS-based discovery techniques lack spiked in standards of known concentrations, making a mathematically based normalization scheme impossible. To explore this issue in the context of discovery proteomics, the normalized relative abundances of several proteins were examined on an individual protein level, then a global strategy was applied to all detected proteins across all samples to test the potential effect of tissue scaling on measured protein abundances.

To investigate the effects of an increased placental tissue mass on protein expression, the normalized relative abundance of each protein from **Figure 4** was multiplied by the normalized relative abundance of PLAP to account for the increased total protein amount as placental tissue mass increases throughout pregnancy. The results are shown in **Supplemental Figure 3.** As shown in the plots, multiplying by PLAP to account for tissue growth abrogates the differential expression trends and instead shows a nearly uniform, statistically significant increase in all but one SLC transporter over gestation (Wilcoxon pairwise testing used to calculate significance between trimesters).

**Supplemental Appendix 1: R Scripts for Proteomic Data Analysis and Figure Generation**

R scripts and packages used to process data after Skyline export and generate figures.

```
#Processing files from placental microsome batches. The single replicate
#injections from each batch were processed individually in Skyline to
#filter on 1 unique peptide/protein with all repeats, duplicates and
#missing removed as well as a 0.8 dotp filter.

library(MSstats)
library(ggplot2)
library(dplyr)
library(tidyverse)
library(PCAtools)
library(limma)
library(tibble)
library(rlang)
library(writexl)
library(openxlsx)
library(viridis)
library(grid)
library(sva)
library(RColorBrewer)

#Import annotation and data files from the desired directory. Used Skyline's
#export feature to export both MS data and annotation files in the appropriate
#MS Stats readable format.

annotb1_term <-read.csv("MSStats Annotation_Batch1_term.csv")

annotb2_term <-read.csv("MSStats Annotation_Batch2_term.csv")

annotb3_term <-read.csv("MSStats Annotation_Batch3_term.csv")

annot_combined_term <-rbind(annotb1_term,annotb2_term,annotb3_term)

batch1 <-read.csv("MSstatsInput4_Batch1_rerun_1pep.csv")
batch2 <-read.csv("MSstatsInput4_Batch2_rerun_1pep.csv")
batch3 <-read.csv("MSstatsInput4_Batch3_rerun_1pep.csv")

batchdata_combined <-rbind(batch1,batch2,batch3)

#Data pre-processing with MS Stats package. Applies normalization, summarization
#and log transformation. No other peptide filtereing is performed at this step
#using MS Stats because of the filtering criteria applied to the data prior
#to Skyline export.

batchdata_import_term <-SkylinetoMSstatsFormat(batchdata_combined,
```

```
                                    annotation = annot_combined_term,
                                    filter_with_Qvalue = FALSE,
                                    useUniquePeptide = FALSE,
                                    removeFewMeasurements = TRUE,
                                    removeProtein_with1Feature = FALSE)


term_q_tmp <-dataProcess(batchdata_import_term,
                                   normalization = "quantile",
                                   summaryMethod = "TMP",
                                   equalFeatureVar = FALSE,
                                   censoredInt = "0",
                                   MBimpute= FALSE,
                                   remove50missing = FALSE)

#Pull out protein level data and transpose matrix for remaining data analysis
#workflow

norm_protein <- term_q_tmp$ProteinLevelData

norm_protein <- subset(norm_protein, select = c(2,3,4))

norm_protein <- norm_protein %>%
  pivot_wider(names_from = originalRUN, values_from = LogIntensities)

#Re-order row names in metadata and column names in protein abundance df to match

align_columns_safe <- function(data_df, metadata, strict = TRUE) {
  stopifnot(!is.null(rownames(metadata)))
  stopifnot(!is.null(colnames(data_df)))

  meta <- rownames(metadata)
  dat  <- colnames(data_df)
  idx  <- match(meta, dat)

  missing <- meta[is.na(idx)]
  extra   <- setdiff(dat, meta)

  if (length(missing)) {
    warning(sprintf("missing in data_df: %s", paste(missing, collapse = ", ")))
  }
  if (length(extra)) {
    message(sprintf("extra columns in data_df: %s", paste(extra, collapse = ", ")))
  }

  if (strict) {
```

```
    if (any(is.na(idx))) {
      message("strict mode: no changes made (incomplete match).")
      return(list(data = data_df, metadata = metadata,
                  matched = FALSE, missing = missing, extra = extra))
    }
    tmp <- data_df[, idx, drop = FALSE]
    if (!identical(colnames(tmp), meta)) {
      warning("strict mode: no changes made (post-order names differ).")
      return(list(data = data_df, metadata = metadata,
                  matched = FALSE, missing = missing, extra = extra))
    }
    return(list(data = tmp, metadata = metadata[meta, , drop = FALSE],
                matched = TRUE, missing = missing, extra = extra))
  } else {
    # common-only
    common <- intersect(meta, dat)
    idx2   <- match(common, dat)
    data_aligned <- data_df[, idx2, drop = FALSE]
    meta_aligned <- metadata[common, , drop = FALSE]
    stopifnot(identical(rownames(meta_aligned), colnames(data_aligned)))
    return(list(data = data_aligned, metadata = meta_aligned,
                matched = length(common) == length(meta),
                missing = setdiff(meta, common), extra = setdiff(dat, common)))
  }
}

align <- align_columns_safe(norm_protein, metadata, strict = TRUE)

norm_protein <- align$data
metadata <- align$metadata

identical(colnames(norm_protein),
          rownames(metadata))

#Proceed to limma for batch effect removal and differential analysis.
#NAs need to be removed

keep_proteins <- rowMeans(!is.na(norm_protein)) >= 1
norm_protein_f <- norm_protein[keep_proteins, ]

metadata$Term <- factor(metadata$Term)
metadata$Batch <- factor(metadata$Batch)


design <- model.matrix(~ 0 + Term + Batch , data = metadata)
```

```
contrast.matrix <- makeContrasts(
  Term2vsTerm1 = Term2 - Term1,
  Term3vsTerm1 = Term3 - Term1,
  Term3vsTerm2 = Term3 - Term2,
   levels = design
)


fit <- lmFit(norm_protein_f, design)
fit2 <- contrasts.fit(fit, contrast.matrix)
fit2 <- treat(fit2, lfc = log2(1), trend = TRUE, robust = TRUE)


topTreat_t2vt1 <- topTreat(fit2, coef = 1, number = Inf)  # Term2 vs Term1
topTreat_t3vt1<- topTreat(fit2, coef = 2, number = Inf)  # Term3 vs Term1
topTreat_t3vt2<- topTreat(fit2, coef = 3, number = Inf)  # Term3 vs Term2

##################FILE AND FIGURE GENERATION##############
#Generate Excel-readable table of resutls from limma output
# Add protein names as a column
topTreat_t2vt1$Protein <- rownames(topTreat_t2vt1)
topTreat_t3vt1$Protein <- rownames(topTreat_t3vt1)
topTreat_t3vt2$Protein <- rownames(topTreat_t3vt2)

# Add flags and labels
add_flags <- function(df, label) {
  df$Comparison <- label
  df$Pass_pval <- df$adj.P.Val < 0.05
  df$Pass_logFC <- abs(df$logFC) > 1
  df$Significant <- df$Pass_pval & df$Pass_logFC
  df$Upregulated <- df$Significant & df$logFC > 1
  df$Downregulated <- df$Significant & df$logFC < -1
  return(df)
}

df_t2t1 <- add_flags(topTreat_t2vt1, "Term2vsTerm1")
df_t3t1 <- add_flags(topTreat_t3vt1, "Term3vsTerm1")
df_t3t2 <- add_flags(topTreat_t3vt2, "Term3vsTerm2")


# Combine all for export
all_proteins <- rbind(df_t2t1, df_t3t1, df_t3t2)

# Create summary table
make_summary <- function(df, label) {
  data.frame(
```

```
    Comparison = label,
    Total_Proteins = nrow(df),
    Pass_adjPval = sum(df$Pass_pval),
    Pass_logFC = sum(df$Pass_logFC),
    Significant = sum(df$Significant),
    Upregulated = sum(df$Upregulated),
    Downregulated = sum(df$Downregulated)
  )
}

summary_df <- do.call(rbind, list(
  make_summary(df_t2t1, "Term2vsTerm1"),
  make_summary(df_t3t1, "Term3vsTerm1"),
  make_summary(df_t3t2, "Term3vsTerm2")
  ))

# Export to Excel with multiple sheets
write.xlsx(
  list(
    AllProteins = all_proteins,
    Summary = summary_df
  ),
  file = "significant_limma_trend.xlsx",
  rowNames = FALSE
)


#Volcano plot function for Figures 1 and 3
make_volcano_plot <- function(results_df, contrast_name, fc_cutoff = 1, p_cutoff = 0.05) {
  # Add regulation column
  results_df$abundance <- with(results_df, ifelse(
    logFC > fc_cutoff & adj.P.Val < p_cutoff, "Higher in T2",
    ifelse(logFC < -fc_cutoff & adj.P.Val < p_cutoff, "Higher in T1", "Not Significant")
  ))

  # Ensure abundance is a factor with all levels
  results_df$abundance <- factor(results_df$abundance, levels = c("Higher in T2",
                                              "Higher in T1", "Not Significant"))

  # Add transformed p-value
  results_df$log10_adjpvalue <- -log10(results_df$adj.P.Val)

  # Plot
  ggplot(results_df, aes(x = logFC, y = log10_adjpvalue, color = abundance)) +
    geom_point(size = 4) +
    scale_color_manual(
```

```
    values = c(
      "Higher in T2" = "#DC3220",
      "Higher in T1" = "#005AB5",
      "Not Significant" = "black"
    ),
    breaks = c("Higher in T2", "Higher in T1", "Not Significant"),  # Only show these in the
legend
    name = NULL
  ) +
  ggrepel::geom_text_repel(aes(label = ifelse(abundance != "Not Significant",
                                              rownames(results_df), "")), size = 6,
                           show.legend = FALSE) +
  geom_vline(xintercept = c(-fc_cutoff, fc_cutoff), col = "blue", linewidth = 1.2,
             linetype = "dashed") +
  geom_hline(yintercept = -log10(p_cutoff), col = "green", linewidth = 1.2) +
  xlab("Log2 Fold Change") +
  ylab("-Log10 Adjusted p-value") +
  ggtitle(paste(contrast_name)) +
  theme_minimal(base_size = 18) +
  theme(
    panel.grid.major = element_blank(),
    panel.grid.minor = element_blank(),
    legend.position = "right",
    axis.line = element_line(color = "black"),
    plot.title = element_text(size = 26),
    axis.line.x = element_line(color = "black", linewidth = 1.5),
    axis.line.y = element_line(color = "black", linewidth = 1.5),
    axis.text = element_text(size = 24),
    axis.title = element_text(size = 22),
    legend.text = element_text(size = 22)
  )
}

vp_limma_t2_t1_trend <- make_volcano_plot(topTreat_t2vt1, "T2/T1 Protein Abundance")
ggsave("volcano_limma_t2vt1_trend.jpg", plot = vp_limma_t2_t1_trend, width = 10,
       height = 8, dpi = 300)


#Batch removal for PCA visualization
#Extract batch information as a factor vector
batch_vector <- as.factor(metadata$Batch)

#Need a separate design matrix without batch

design_nb <- model.matrix(~ 0 + Term, data = metadata)
```

```
norm_protein_bc <- removeBatchEffect(norm_protein_f, batch = batch_vector,
                                     design = design_nb )


#Process MS Stats transformed output using "PCAtools"

p_limma <- pca(combined_limmabc, metadata = metadata, removeVar = 0.1)

#elbow method for PCA inclusion

horn <- parallelPCA(combined_limmabc)
horn$n

elbow <- findElbowPoint(p_limma$variance)
elbow

which(cumsum(p_limma$variance) > 80)[1]


screeplot(p_limma,
          components = getComponents(p_limma, 1:20),
          vline = c(horn$n, elbow)) +

  geom_label(aes(x = horn$n + 1, y = 50,
             label = 'Horn\'s', vjust = -2, hjust = 1.5, size = 8)) +
  geom_label(aes(x = elbow + 1, y = 50,
             label = 'Elbow method', vjust = -2, size = 8))

#Use to generate Figure 2D

p_limma$Term <- as.numeric(p_limma$Term)
p_msstats$Batch <- as.numeric(as.factor(p_msstats$Batch))
p_limma$Sex <- as.numeric(as.factor(p_limma$Sex))

p_limma$metadata$Term <- as.factor(p_limma$metadata$Term)
p_limma$metadata$Batch <- as.factor(p_limma$metadata$Batch)
p_limma$metadata$Sex <- as.factor(p_limma$metadata$Sex)

pca_biplot <- biplot(p_limma, x = "PC1", y = "PC2",
            showLoadings = TRUE,
            lengthLoadingsArrowsFactor = 1.5,
            sizeLoadingsNames = 4,
            colLoadingsNames = 'black',
            lab = NULL,
            labSize = 6,
            colkey = c('1' = "blue", '2'= "green", '3' = "orange", p_limma$metadata$Term),
```

```
          colby = 'Term',
          hline = 0, vline = 0,
          vlineType = 'solid',
          hlineType = 'solid',
          gridlines.major = FALSE, gridlines.minor = FALSE,
          pointSize = 5,
          legendPosition = 'left', legendLabSize = 14, legendIconSize = 8.0,
          #shape = 'Batch', shapekey = c('1'=15, '2'=17, '3'=8),
          drawConnectors = FALSE,
          #title = 'PCA',
          subtitle = 'PC1 versus PC2')

# Create a named vector for custom colors
group_levels <- levels(p_limma$metadata$Term)
custom_colors <- setNames(c("blue", "green", "orange"), group_levels)

ggsave("test.jpeg", plot = pca_biplot, device = "jpeg",
      width = 8, height = 5.5, dpi = 3000)


#Protein abundance boxplots with limma/treat star labels used to generate Figures 4, 5 and
Supplemental Figures 1 and 3

plot_protein_box_limma <- function(
   norm_mat_bc,          # matrix/data.frame: rows = proteins, cols = samples (batch-removed)
   metadata,             # data.frame: rownames = sample IDs; must contain column 'Term' (1/2/3
or T1/T2/T3)
   all_proteins,         # data.frame: limma/treat results with columns: Protein, Comparison,
adj.P.Val (logFC optional)
   protein_id,           # character: protein to plot (must be a rowname in norm_mat_bc)
   protein_full_name = NULL,  # optional: full protein name for title
   term_levels = c("T1","T2","T3"),
   palette = c("T1"="#0072B2","T2"="#E69F00","T3"="#CC79A7"),
   title = NULL,
   y_label = "Normalized Intensity",
   y_pad_frac = 0.07,
   pair_to_comp_fn = NULL   # optional: function mapping c("T1","T2") -> "Term2vsTerm1"
) {
  if (is.null(pair_to_comp_fn)) {
   pair_to_comp_fn <- function(pair) {
     stopifnot(length(pair) == 2)
     a <- sub("^T", "", pair[1])  # left (group1)
     b <- sub("^T", "", pair[2])  # right (group2)
     paste0("Term", b, "vsTerm", a)
   }
  }
```

```
stars_from_p <- function(p) {
  ifelse(p <= 1e-4, "****",
         ifelse(p <= 1e-3, "***",
                ifelse(p <= 1e-2, "**",
                       ifelse(p <= 5e-2, "*", "ns"))))
}


if (is.data.frame(norm_mat_bc)) norm_mat_bc <- as.matrix(norm_mat_bc)
if (is.null(rownames(norm_mat_bc))) stop("norm_mat_bc must have rownames = protein IDs.")
if (!protein_id %in% rownames(norm_mat_bc)) stop(sprintf("Protein '%s' not found in
norm_mat_bc.", protein_id))
if (!("Term" %in% colnames(metadata))) stop("metadata must contain a 'Term' column.")
if (!identical(rownames(metadata), colnames(norm_mat_bc))) {
  stop("Sample alignment mismatch: rownames(metadata) must exactly equal
colnames(norm_mat_bc).")
}
req_cols <- c("Protein","Comparison","adj.P.Val")
if (!all(req_cols %in% colnames(all_proteins))) {
  stop(sprintf("all_proteins must contain columns: %s", paste(req_cols, collapse = ", ")))
}

sample_ids <- colnames(norm_mat_bc)
subset_df <- data.frame(
  Sample    = sample_ids,
  intensity = as.numeric(norm_mat_bc[protein_id, sample_ids]),
  Term_raw  = metadata[sample_ids, "Term"],
  stringsAsFactors = FALSE
)

# Map Term
term_map <- setNames(term_levels, c("1","2","3"))
term_char <- as.character(subset_df$Term_raw)
term_mapped <- ifelse(term_char %in% names(term_map), term_map[term_char], term_char)
subset_df$Term <- factor(term_mapped, levels = term_levels)
subset_df$Term_raw <- NULL

# ----- build annotation df from limma/treat results -----
lev <- levels(droplevels(subset_df$Term))
pairs <- if (length(lev) >= 2) utils::combn(lev, 2, simplify = FALSE) else list()

y_max <- max(subset_df$intensity, na.rm = TRUE)
y_min <- min(subset_df$intensity, na.rm = TRUE)
y_range <- y_max - y_min
if (!is.finite(y_range) || y_range == 0) y_range <- 1
y_step <- y_pad_frac * y_range
```

```
  ann_list <- vector("list", length(pairs))
  for (i in seq_along(pairs)) {
    pair <- pairs[[i]]
    comp <- pair_to_comp_fn(pair)
    rows <- all_proteins[all_proteins$Protein == protein_id & all_proteins$Comparison == comp,
]
    if (nrow(rows) == 0) {
      ann_list[[i]] <- data.frame(
        group1     = pair[1],
        group2     = pair[2],
        p.adj      = NA_real_,
        label      = NA_character_,
        y.position = y_max + i * y_step,
        stringsAsFactors = FALSE
      )
    } else {

      pval <- as.numeric(rows$adj.P.Val[1])
      raw_label <- stars_from_p(pval)

      ann_list[[i]] <- data.frame(
        group1     = pair[1],
        group2     = pair[2],
        p.adj      = pval,

#label      = raw_label,
        label      = ifelse(raw_label == "ns", NA, raw_label), #(removes ns from plots)
        y.position = y_max + i * y_step,
        stringsAsFactors = FALSE
      )

    }
  }

  ann_df <- if (length(ann_list) > 0) {
    do.call(rbind, ann_list)
  } else {
    data.frame(
      group1 = character(),
      group2 = character(),
      p.adj = numeric(),
      label = character(),
      y.position = numeric(),
      stringsAsFactors = FALSE
    )
```

```
  }

  # ----- plot -----
  title_text <- if (is.null(title)) {
    if (!is.null(protein_full_name)) {
      paste0(protein_id, " (", protein_full_name, ")")
    } else {
      protein_id
    }
  } else {
    title
  }
  p <- ggplot2::ggplot(subset_df, ggplot2::aes(x = Term, y = intensity, fill = Term)) +
    ggplot2::geom_boxplot(outlier.shape = 21, outlier.size = 2, alpha = 0.9) +
    ggplot2::labs(title = title_text, x = "Term", y = y_label) +
    ggplot2::scale_fill_manual(values = palette) +
    ggplot2::theme_minimal() +
    ggplot2::theme(
      panel.grid  = ggplot2::element_blank(),
      axis.line   = ggplot2::element_line(color = "black", linewidth = 1.8),
      axis.text   = ggplot2::element_text(size = 22),
      axis.title  = ggplot2::element_text(size = 22),
      plot.title  = ggplot2::element_text(size = 22, face = "bold")
    )+
    ggplot2::theme(legend.position = "none")

  if (nrow(ann_df) > 0 && all(c("group1","group2","label","y.position") %in%
colnames(ann_df))) {

    p <- p + ggpubr::stat_pvalue_manual(
      ann_df,
      label       = "label",
      y.position  = "y.position",
      xmin        = "group1",
      xmax        = "group2",
      tip.length  = 0.01,
      size        = 8,     # <- font size for stars/"ns"
      bracket.size= 1.0,   # <- thickness of the bracket lines
      # optional extras:
      # step.increase = 0.06,  # vertical spacing between multiple brackets
      # vjust = -0.2,          # label vertical position relative to bracket
      # label.size = 0.0       # outline around text; 0 removes stroke
    )

  }
```

```
  return(p)
}

ggsave("meanplot_term.png", plot = last_plot(), width = 8, height = 6,
       dpi = 300)
```

### Supplemental Figures

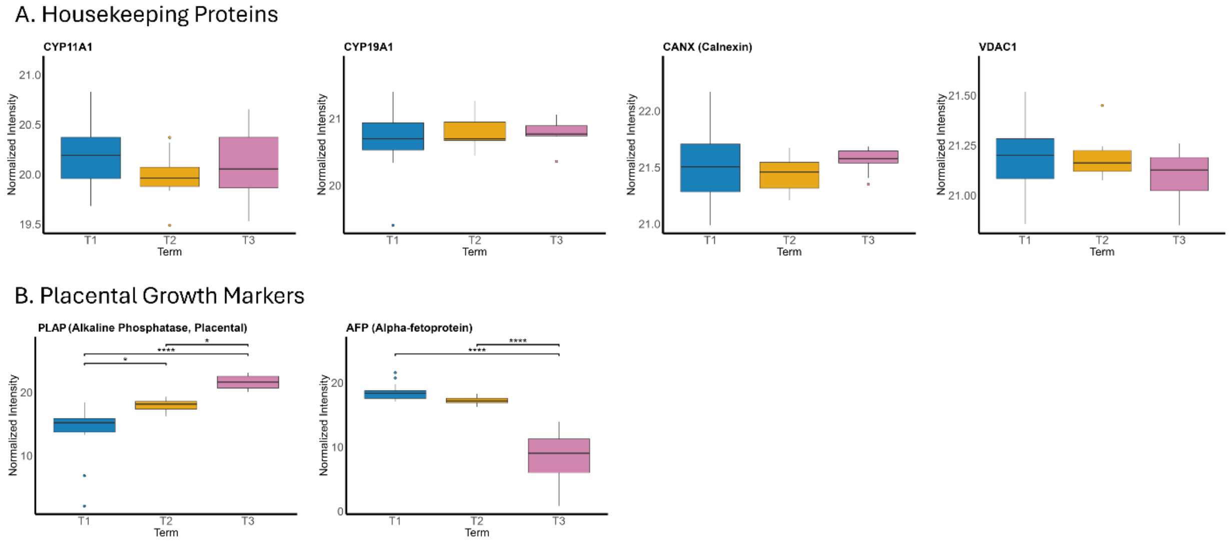


**Supplemental Figure 1.** Individual plots comparing protein abundance across trimesters for select housekeeping proteins (A) and two placental growth markers (B). Housekeeping proteins show no significant variation between gestational ages. PLAP shows an increase in relative abundance with increasing gestational age while AFP shows a decrease. Black bars represent the median value of the normalized, log-transformed, batch corrected abundance value per trimester. Boxes denote the interquartile range between 25% and 75% of the normalized intensity range. Whiskers extend to cover 1.5x the interquartile range and dots represent outlier samples. Statistical significance was computed using the Benjamini-Hochberg method applied to the raw *p*-values calculated from empirical Bayes analysis after linear modeling trends in the data using trimester comparisons as contrasts. * = $p < 0.05$, ** = $p < 0.01$, *** = $p < 0.001$, and **** = $p < 0.0001$.

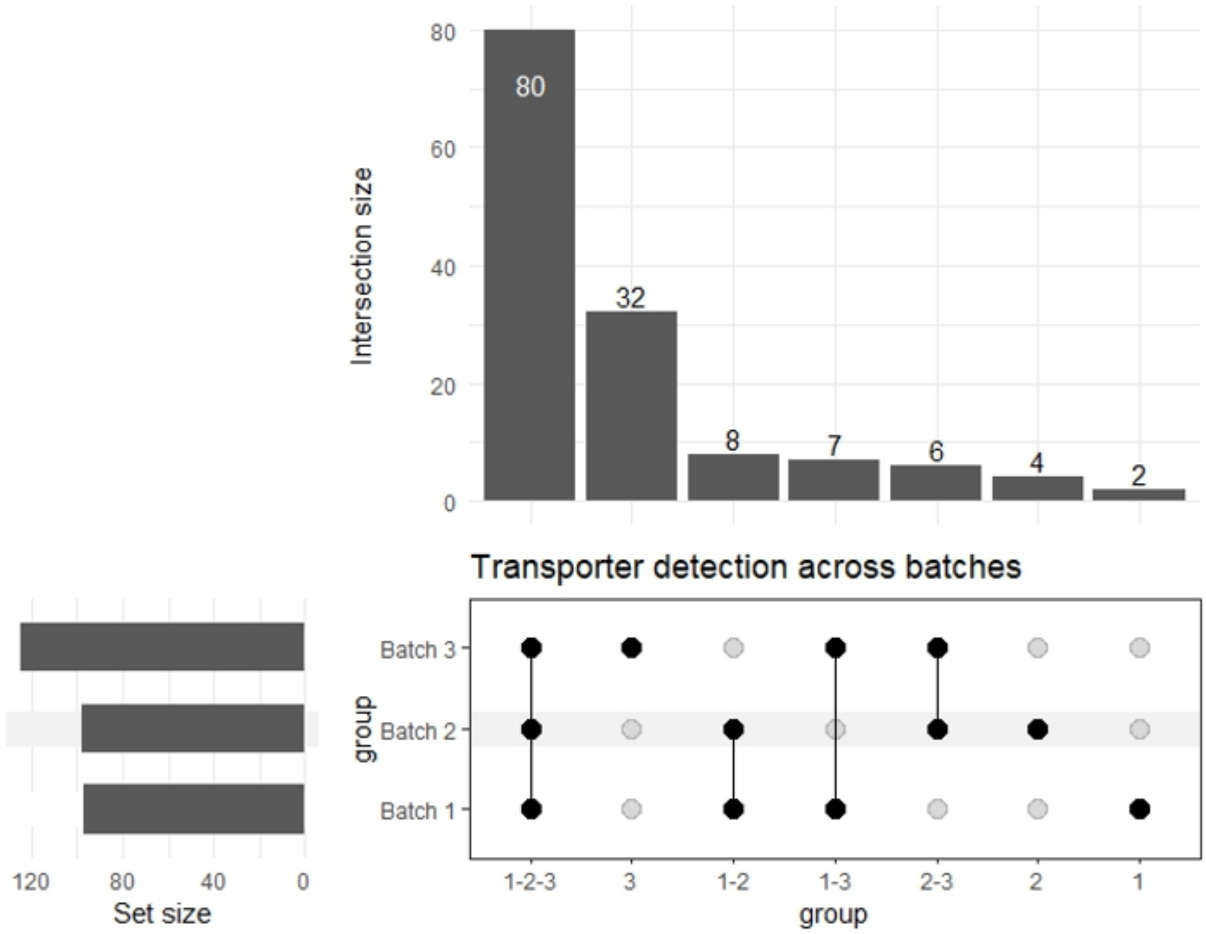


**Supplemental Figure 2.** Upset plot used to track overlap and differences between transport protein detections across DIA batches. Batches 1 and 2 contain similar numbers of transport proteins ($n$ = 97 and $n$ = 98, respectively) with minimal overlap while batch 3 contains the highest number of transporter IDs ($n$ = 125). Overall, 80 transporters were shared across all three batches, 32 were unique to batch 3, four were unique to batch 2 and two were unique to batch 1.

**Supplemental Figure 3.** Individual plots after scaling for total placental growth using PLAP.

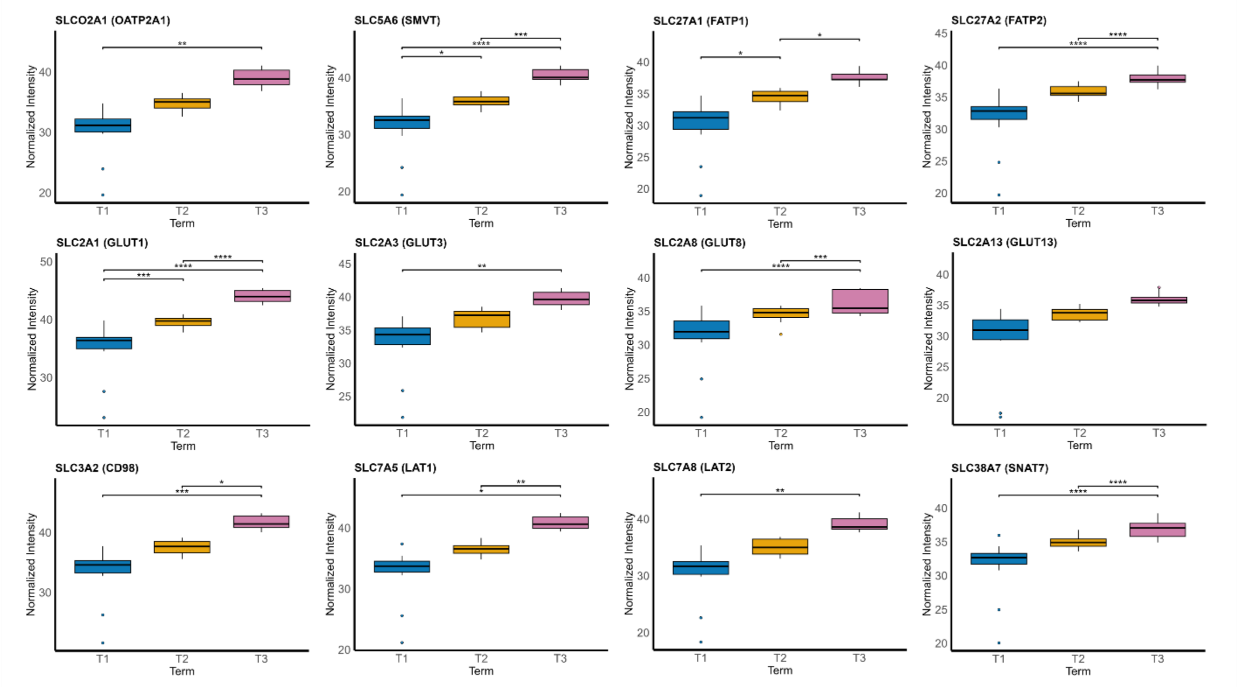


Normalized abundance values for individual transporters were multiplied by the normalized abundance value for PLAP in each sample. Black bars represent the median value of the normalized, log-transformed, batch corrected abundance value per trimester. Boxes denote the interquartile range between 25% and 75% of the normalized intensity range. Whiskers extend to cover 1.5x the interquartile range and dots represent outlier samples. Statistical significance was determined using Wilcoxon pairwise testing between trimesters. * = $p < 0.05$, ** = $p < 0.01$, *** = $p < 0.001$, and **** = $p < 0.0001$.
